# Choice history biases evidence accumulation: a cross-species comparison from humans to mice

**DOI:** 10.1101/2025.08.22.671879

**Authors:** Kianté Fernandez, Alexander Fengler, Anne E. Urai

## Abstract

Mice are increasingly used to study the neural circuit-level basis of behavior, often with the ultimate goal of extrapolating these insights to humans. To generalize insights about neural functioning across species, it is crucial to ensure correspondence in behavioral and cognitive strategy. We previously showed that human observers’ evidence accumulation is biased by previous choices (***Urai et al., 2019***). To replicate these findings across species, we fit Diffusion Decision Models (***Fengler et al., 2025***) to behavioral data from 62 mice performing a standardized perceptual decision-making task (***The International Brain Laboratory et al., 2021***). We identified the same cognitive strategy of history-dependent evidence accumulation: individual differences in choice repetition were explained by a history-dependent bias in the rate of evidence accumulation rather than its starting point. We argue that history-biased evidence integration reflects a fundamental aspect of perceptual decision-making, that may transcend the specific species.

## Introduction

In perceptual decision-making, evidence accumulation is biased by previous choices: our previous work has shown that in human this seems to happen through a history-dependent change in the speed of evidence accumulation (drift bias), rather than in via a shift starting point of the accumulation process (***Urai et al., 2019***) (Figure 1). Choice history (defined as one or more past choices) thus seems to bias the interpretation of current sensory input, akin to shifting endogenous attention toward (or away from) the previously selected interpretation of sensory evidence (***Urai and Donner, 2022***). This decision-making strategy, which robustly captures individual differences across tasks, may be exhibited across mammalian species—thereby providing insights into the shared neural circuit mechanisms of cognition.

**Figure 1.**
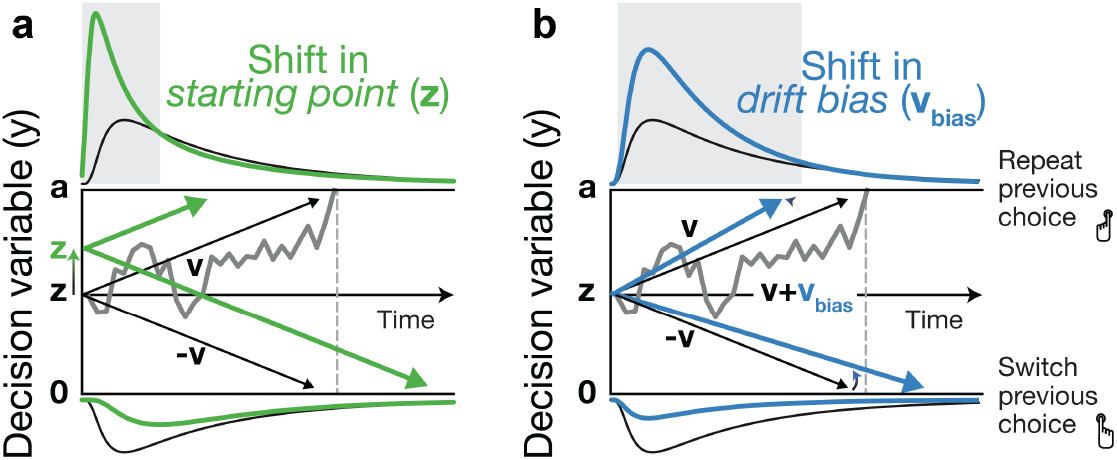
The DDM postulates that noisy sensory evidence is accumulated over time, until the resulting decision variable *y* reaches one of two bounds (lines at *y* = 0 and *y* = *a*) for the two choice options. Repeating this process over many trials yields RT distributions for each choice option respectively plotted above and below the bounds. Gray line: example trajectory of the decision variable from a single trial. Black lines: mean drift and resulting RT distributions under unbiased conditions. **(a)** Choice history-dependent shift in starting point. Green lines: mean drift and RT distributions under biased starting point. Gray-shaded area indicates those RTs for which starting point leads to choice bias. **(b)** Choice history-dependent shift in drift bias. Blue lines: mean drift and RT distributions under biased drift. Gray shaded area indicates those RTs for which drift bias leads to choice bias. Adapted under a CC-BY license from (***Urai et al., 2019***).

Recent advances in training mice to perform complex tasks, combined with powerful neural measurement tools, have positioned mice as a popular model species in cognitive neuroscience research (***Urai et al., 2022***). An important premise is that cognitive and computational mechanisms of behavior are preserved across species. However, this is rarely explicitly tested, creating challenges in the ability to translate mice-based neuroscience findings to humans (***Barron et al., 2020***). Here, we directly replicate our previous analyses in humans (***Urai et al., 2019***) in mice. We analyzed choices and response times of 62 mice performing a standardized decision task (***The International Brain Laboratory et al., 2021***) and show that animals exhibit the same computational strategy as humans: a history-dependent drift bias.

## Results

Choice repetition in perceptual tasks has been reported across species from rodents to monkeys and humans (***Busse et al., 2011***; ***Gold et al., 2008***; ***Fründ et al., 2014***). The basis of this work is an open dataset from the International Brain Laboratory (***The International Brain Laboratory et al., 2021***), where animals are trained to detect the location of a visual stimulus on a screen, which varies in contrast (Figure 2a). Animals then use a wheel to move the stimulus into the center of the screen to obtain a water reward.

**Figure 2.**
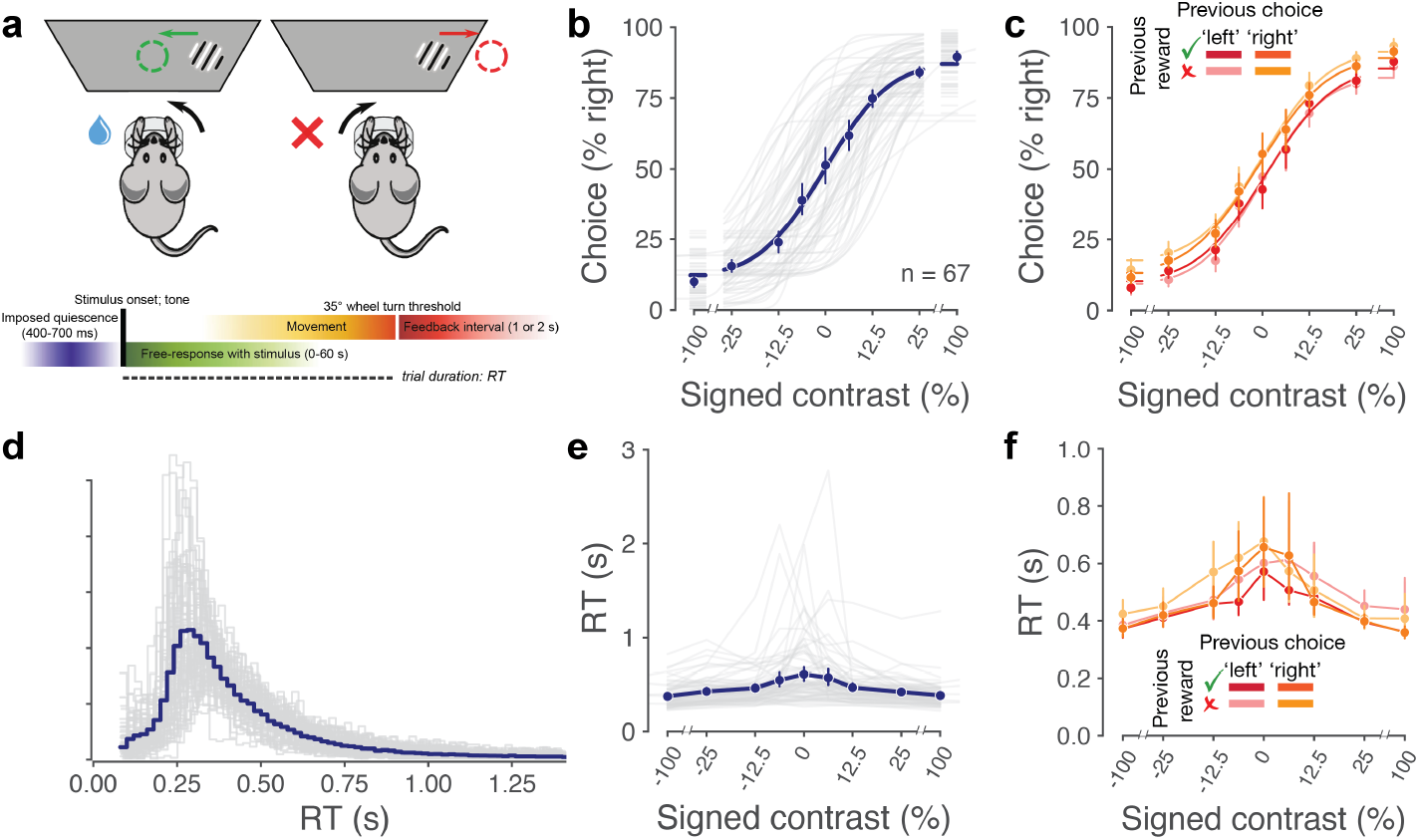
Mice repeat their choices. **(a)** A standardized decision-making task, adapted from (***The International Brain Laboratory et al., 2021***) under a CC-BY license. **(b)** Psychometric functions for all animals (each shown with a grey line). **(c)** History-conditioned psychometric functions. **(d)** Distributions of response times, taken as the time from stimulus onset until response completion: either bringing the stimulus into the center of the screen, or the equivalent distance off the screen. Each animal is shown with one grey line. **(e)** Chronometric functions, showing median RT per contrast level for all animals (each shown with a grey line). **(f)** History-conditioned chronometric functions.

We here selected data from early in the animals’ training: after animals had sufficiently learned the basic task, but before the introduction of a block manipulation that complicates analyses of choice repetition. As shown in previous analyses of these data (***The International Brain Laboratory et al., 2021***; ***Findling et al., 2025***), mice tend to repeat their past choice irrespective of the past trial’s outcome (correct or error) (Figure 2).

### Fitting history-dependent sequential sampling models

To distinguish different mechanisms that may underlie this repetition bias, we fit four variants of a Diffusion-Decision Model (***Ratcliff et al., 2016***): with choice history affecting no parameters, only *z*, only *v*_*bias*_, or both *z* and *v*_*bias*_. Model comparison favors models where only *v*_*bias*_ or both parameters scale with choice history, replicating our previous results in humans (Figure 3a).

**Figure 3.**
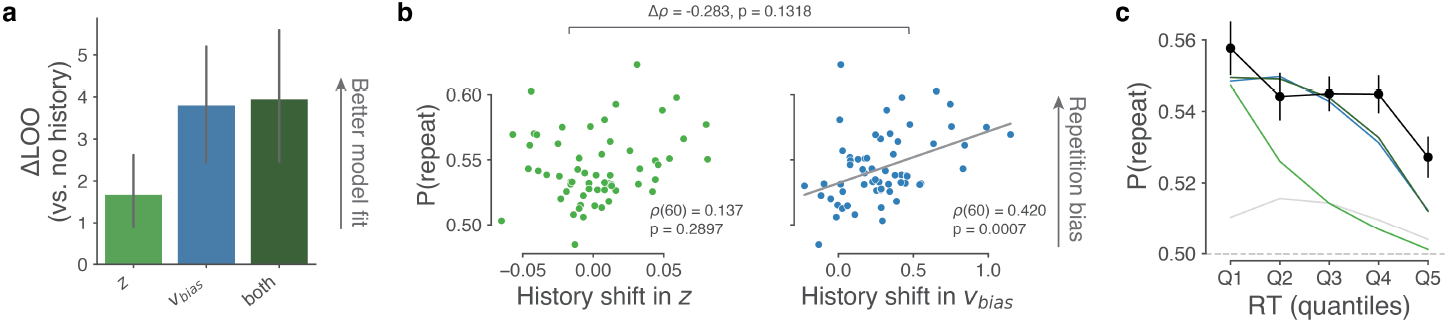
Individual choice history biases are explained by history-dependent changes in drift bias, not starting point. **(a)**. Model comparison of models where *z, v*_*bias*_ or both were allowed to vary as a function of the animals’ previous choice. The LOO for a model without history dependence was used as a baseline for each dataset. Higher ΔLOO values indicate a model that is better able to explain the data, taking into account the model complexity. **(b)** Correlation between individual choice repetition probabilities, P(repeat), and history shift in starting point (left column, green) and drift bias (right column, blue). Parameter estimates were obtained from a model in which both bias terms were allowed to vary with previous choice. Grey line shows the best fit of a linear regression, only plotted for Spearman’s *ρ* correlations significant at p<0.05. Top: Δ*ρ* value between the two correlation coefficients, with p-value computed from a Steiger’s test. **(c)** Conditional bias functions comparing observed data with simulations from four DDM variants. The probability of repeating a previous choice, P(repeat), is plotted across five reaction time (RT) quantiles. The observed data (black) show that choice repetition is most frequent on fast trials and decreases as RT increases. This empirical pattern is captured by the best-fitting *v*_*bias*_ model (blue).

The results from quantitative model comparisons were corroborated by inspection of individual differences. Animals’ choice repetition probability correlated with their history shift in *v*_*bias*_, but not in *z* (Figure 3b). Conditional bias functions confirm this qualitatively: animals show repetition bias across all RT quantiles, a pattern expected from a *v*_*bias*_ but not *z* (Figure 3). This replicates our previous finding in six human perceptual decision-making datasets (***Urai et al., 2019***).

## Discussion

In this work, we test various formulations of sequential sampling models (***Fengler et al., 2022***) to a public dataset of mouse decision-making (***The International Brain Laboratory et al., 2021***) and find that mice display the same history-dependent drift biases as previously shown in humans (***Urai et al., 2019***). Evidence accumulation over multiple temporal scales may thus reflect a fundamental aspect of decision-making, conserved across mammalian species.

These findings set the stage for linking the computations of decision-making to neural dynamics at the single-cell and population levels. Specifically, recent work showed that an exponentially weighted average of past choices informs an animal’s prior, which can be decoded across brain regions (***Findling et al., 2025***). Future work could extend these findings by developing neurocognitive models that link neural activity at both the single-cell and population levels with behavioral modalities through integrative modeling (***Nunez et al., 2024***). Such joint modeling can enable the simultaneous characterization of single-trial neural measures and cognitive modeling parameters, thereby capturing trial-by-trial variations in drift bias (***Ghaderi-Kangavari et al., 2023***).

Our replication of previous human findings has several limitations that we plan to address in future work. We used animals’ single past choice at *t* − 1 as a limited read-out for the full prior that is constructed from past evidence: future work could include choice history beyond one trial (***Fründ et al., 2014***; ***Findling et al., 2025***; ***Urai et al., 2019***). Moreover, choice history bias may arise both from true repetition behavior and from slow drifts in decision criterion (***Gupta and Brody, 2022***; ***Vloeberghs et al., 2025***; ***Urai, 2025***), which may in turn affect trial-to-trial fluctuations in drift bias. Lastly, mice are known to occupy different engagement states over the course of a session (***Ashwood et al., 2022***; ***Urai et al., 2019***), which may need to be estimated to better fit sequential sampling models (***Chakravarty et al., 2025***). All of these extensions may ultimately allow a richer estimation of the interaction between latent computational variables that vary across and within trials (***Urai, 2025***).

## Methods and Materials

### Task, procedure and data selection

Mouse data were collected by the International Brain Laboratory (***The International Brain Laboratory et al., 2021***), and released as part of (***Bruijns et al., 2023***). Mice were trained to perform a perceptual decision-making task, reporting the location of a visual grating (of varying contrast) by turning a steering wheel left or right. We selected sessions where mice had mastered the basic task but had not yet been exposed to structured stimulus sequences with biased probabilities. This approach ensured that behavioral choice history biases reflected endogenous, rather than experimentally induced, biases. For each mouse, we analyzed three sessions just prior to achieving training proficiency in the basic task, ensuring comparable performance levels across animals while avoiding any influence of the probabilistic stimulus sequences introduced in later training stages.

We defined RT as the difference between stimulus onset and completion of the response. Note that this differs slightly from previous work on this dataset (***The International Brain Laboratory et al., 2021***), which used the time of first movement initiation rather than choice completion as reaction time. While likely a mixture of multiple processes, we chose to include the entire interval as our measure of the decision process, as we aim to evaluate the DDMs capacity to capture components of processing associated with the entire period the animal engaged in the task.

We included sessions based on several quality control criteria: each session included at least five contrast levels; performance above 80% correct on high-contrast trials (≥50% contrast); a median RT of less than 1 second; no negative RTs (likely reflecting hardware failure); and a total of >= 200 trials per session remaining after these exclusion criteria. We included only trials with RTs and movement onset times between 0.08 and 2 seconds - excluding outliers that likely represented anticipatory or disengaged responses (***The International Brain Laboratory et al., 2025***). After exclusions, our final dataset contained choices and RTs from 62 animals.

### Cognitive Modeling

To model the choice and RTs of the animals, we use the sequential sampling modeling (SSM) framework (***Forstmann et al., 2016***; ***Fengler et al., 2022***). This widely used class of cognitive models describes decision-making as a stochastic, dynamic evidence accumulation process which evolves one or multiple momentary evidence-states, until a particular predetermined threshold level is reached which triggers a given choice (***Ratcliff and McKoon, 2008***; ***Ratcliff, 1978***). Such “boundary-crossings” trigger a response, leading to distributions of RTs. SSMs thus provide a generative model for reaction time distributions of actions made by organisms, capturing both the speed of the response and the choice itself. Since similar models were used to capture choice history effects in previous work, we only briefly reintroduce the models here and refer readers to ***Urai et al. (2019***) for further details.

#### DDM

In this study we use the basic Diffusion-Decision Model (***Ratcliff et al., 2016***) (also known as Drift Diffusion Model; Figure 1).

The DDM describes the accumulation of noisy sensory evidence which can be described via the following stochastic differential equation:

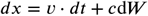

where *x* is the decision variable and *cdW* is Gaussian distributed white noise with mean 0 and variance *c*^2^*dt*. This represents a basic continuous random walk with drift (***Smith, 2000***). The evidence accumulation process terminates when *x* reaches either the upper bound *a* or the lower bound 0 (sometimes this is instead represented as hitting *a* or −*a* instead).

In our work, we represent the drift *v*_*t*_, where *t* refers to a given trial number, via the following simple linear regression:

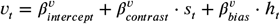

where *s*_*t*_ represents the signed stimulus contrast on trial *t* (negative for left stimuli, positive for right stimuli). Since we observed a saturation of drift rates for contrasts higher than 25%, we clipped all contrast levels above this value to 25% (or −25% for stimuli on the left). *h*_*t*_ is defined as the signed choice (−1 for leftwards, +1 for rightwards) on trial *t* − 1.

We use a similar definition for the starting point:

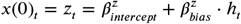

where 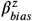 represents how previous choices affect the starting point—a history-dependent bias in the decision process. We compare these against models without history-dependence where we either drop the regression aspect on *v* and/or the regression aspect on *z*, in which case they act as parameters without trial-level adjustments.

Together, 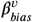 and 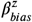 capture how choice history influences decisions through distinct mechanisms. As previously shown, the two biasing mechanisms result in the same (asymmetric) fraction of choices, but they differ in terms of the resulting shapes of RT distributions (Figure 1).

#### Model comparison

To compare model fits across model variants, we estimated out-of-sample predictive performance using LOO cross-validation (***Vehtari et al., 2017***). This approximation, based on Pareto-smoothed importance sampling (PSIS), uses the expected log pointwise predictive density (ELPD) estimates to assess how well a model is likely to fit holdout data without requiring iterative refitting (***Vehtari et al., 2017***). We used the routines implemented in the ArviZ Python package (***Kumar et al., 2019***).

#### Parameter estimation

All models were fit using HSSM (Hierarchical Sequential Sampling Modeling) (***Fengler et al., 2025***), a Python toolbox that facilitates Bayesian inference for a broad class of Sequential Sampling Models, including hierarchical mixed-effects regressions on each underlying cognitive model parameter. To do so, HSSM constructs specified regressions with the Bambi library (***Capretto et al., 2022***) and complies the models with the PyMC probabilistic programming library (***Abril-Pla et al., 2023***). Parameter posteriors and posterior predictive checks were plotted with ArviZ (***Kumar et al., 2019***).

## Data and code availability

All code is available at https://github.com/kiante-fernandez/2023_choicehistory_HSSM.

## Acknowledgments

We thank the International Brain Laboratory for data collection, curation and software development. Philippa Johnson, Charline Tessereau, Naureen Ghani, Naoki Hiratani, Miles Wells, Peter Latham and Alex Pouget provided valuable discussions on the definition of RT in the IBL task. We thank Zeynep Gunes Ozkan for discussions and help in an earlier stage of the project. AEU is supported by a Veni grant (VI.Veni.212.184) from the Netherlands Organization for Scientific Research.

